# Quantitative dynamics of intracellular NMN by genetically encoded biosensor

**DOI:** 10.1101/2023.10.23.563573

**Authors:** Liuqing Chen, Pei Wang, Guan Huang, Wenxiang Cheng, Kaijing Liu, Qiuliyang Yu

**Affiliations:** Sino-European Center of Biomedicine and Health, Institute of Biomedicine and Biotechnology, Shenzhen Institute of Advanced Technology, Chinese Academy of Sciences, 518055 Shenzhen, China; Translational Medicine R&D Center, Institute of Biomedical and Health Engineering, Shenzhen Institutes of Advanced Technology, Chinese Academy of Sciences, 518055 Shenzhen, China; Department of Medical Oncology, Sun Yat-sen University Cancer Center, 510060 Guangzhou, China

## Abstract

Nicotinamide mononucleotide (NMN) is a major precursor for NAD^+^ metabolism with promising effects in treating NAD^+^- and aging-related pathologies. However, measuring live cell NMN dynamics was not possible, leaving key questions in intracellular NMN uptake and regulation unanswered. Here we developed a genetically encoded bioluminescent sensor to quantify subcellular NMN in live cells by fusing engineered NMN-responsive binding domain to bioluminescent and fluorescent proteins from BRET pairs. The sensor dissected the multimechanistic uptake of extracellular NMN and precursors in live cells. We then captured the notably low mitochondrial NMN content and the thereafter vulnerable NMN/NAD^+^ ratio and SARM1 activation in mitochondria, establishing NMN/NAD^+^ ratio as an important parameter in evaluating NAD^+^ boosting strategies. Moreover, we characterized the signature of major NAD^+^ regulating enzymes on NMN and NMN/NAD^+^ ratios, in which Slc25a45 was identified to be a potential mitochondrial NMN transporter for its unique fingerprint on mitochondrial NMN/NAD^+^ ratio.

## INTRODUCTION

Nicotinamide adenine dinucleotide (NAD^+^) regulates numerous biological processes because of its pivotal roles in redox reactions, energy metabolism, as well as gene and protein modifications. Nicotinamide mononucleotide (NMN) is the direct precursor of NAD^+^ in the salvage pathway of NAD^+^ biosynthesis and has been shown to potently boost NAD^+^ in both animal and clinical studies, leading to its potential application in treating a range of NAD^+^-related pathologies^1,2^ such as diabetes,^3^ cardiovascular diseases,^4,5^ Alzheimer’s disease^6^ and skeletal muscle function decline.^7^

However, recent studies suggested NMN may exert regulatory functions beyond a substrate for synthesizing NAD^+^. As NMN and NAD^+^ can compete for the same binding sites in certain enzymes^8^ due to structural similarities, the perturbation of NMN homeostasis in live cells could lead to more profound and widespread regulations. To date, a comprehensive understanding of NMN metabolism and dynamic regulation remains incomplete, particularly regarding NMN’s cellular uptake,^9^ subcellular concentrations^10,11^ and intracellular transport mechanisms.^12^ To address these questions, the precise measurement of NMN levels with spatial and temporal resolution is much needed. Currently, NMN quantification relies on chromatography,^13,14^ mass spectrometry,^15^ enzymatic assays^16,17^ or chemical probes,^18^ all of which require sample lysis and thus compromise the intrinsically rich spatiotemporal information of the biological processes. Genetically encodable biosensors offer a promising alternative,^19,20^ allowing targeted localization in specific subcellular compartments with temporally controlled expression. Several genetically encoded biosensors have been developed for other key metabolites,^21^ but a biosensor for NMN remains notably absent, leaving key questions in NMN and NAD^+^ metabolism unanswered.

Here, we introduce the first genetical encoded nicotinamide mononucleotide ratiometric indicator (NMoRI) capable of quantifying free NMN in both live cells and *in vitro* environments. Using the sensor, we quantified the effects of precursors and various enzymes in NMN homeostasis, elucidating the NMN uptake dynamics and regulations in live cells. Furthermore, NMoRI aided the identification of a potential mitochondrial NMN transporter, extending our understanding of NMN’s intracellular dynamics.

## RESULTS

### Design of NMoRI for sensing cellular NMN

NMoRI is an NMN-sensitive protein domain with its N and C terminus fused respectively to a luciferase and a red fluorescent protein that form a BRET pair. The NMN-sensitive protein undergoes large conformational shift upon binding to NMN and hence affects the relative distance and orientation between the luciferase and the red fluorescent protein, resulting in an NMN-dependent change of the NMoRI’s intramolecular BRET efficiency, which was measured to indicate the NMN concentration of the milieu.

As no NMN-binding protein with desirable conformational changes is available in the existing protein database, we designed the NMN-specific binding domain based on a previously developed protein scaffold for sensing NAD^+^ derived from a catalytically inactive variant of DNA ligase from *Enterococcus faecalis* V583 (*Ef*LigA).^22,23^ Since the NMN molecule contains a negatively charged phosphate group capable of electrostatic interactions with amino acid residues (Figure S1A), we postulated that positively charged residues in the Domain B (K79-P317) of *Ef*LigA could interact with NMN’s phosphate group and thus could pull the Domain A (M1-E78) and B (K79-P317) close (Figure S1B). After testing mutation D91R and Y227A, we obtained the first version of NMN sensor, NMoRI^0.1^ (Figure S1C, D).

NMoRI^0.1^ exhibited a high affinity towards NMN (c50 = 1.34 μM). However, it also responded to NAD^+^ with a c50 of 28.56 μM (Figure S1E). To improve the sensor’s specificity, we then mutated the residues in the NAD^+^ binding pocket from Domain B (S26D, D91R, A125D and Y227K) and obtained NMoRI^0.2^ (Figure S1D, F) with a c50 of 10 μM for NMN and a much-reduced response towards NAD^+^ and other analogues (Figure S1G). Despite of the improvement, NAD^+^ at higher than 100 μM concentrations could still interfere with the NMN measurement. To make NMoRI^0.2^ suitable for cellular applications, where the free intracellular NAD^+^ ranges between 50-300 μM, we further introduced mutation L89M and T228Q to obtain NMoRI^1.0^ (Figure 1A and Figure S1D) with a c50 of 8.0 μM for NMN (Figure 1B and C). NMoRI^1.0^ is NMN-specific and exhibited no significant responses to other analogues within their physiological concentration ranges (Figure 1D). Up to 500 μM of NAD^+^ did not significantly alter NMoRI^1.0^’s response curve (Figure 1E). NMoRI^1.0^ displayed a slight sensitivity to pH changes (Figure S1H). Moreover, NMoRI^1.0^ is not responsive towards high levels (> 1 mM) of AXP (AMP, ADP and ATP) (Figure 1F), but their presence affected the response curves of NMoRI^1.0^ for NMN (Figure S1I, J). NMoRI^1.0^ is hence used in all following experiments and referred to as NMoRI.

**Figure 1.**
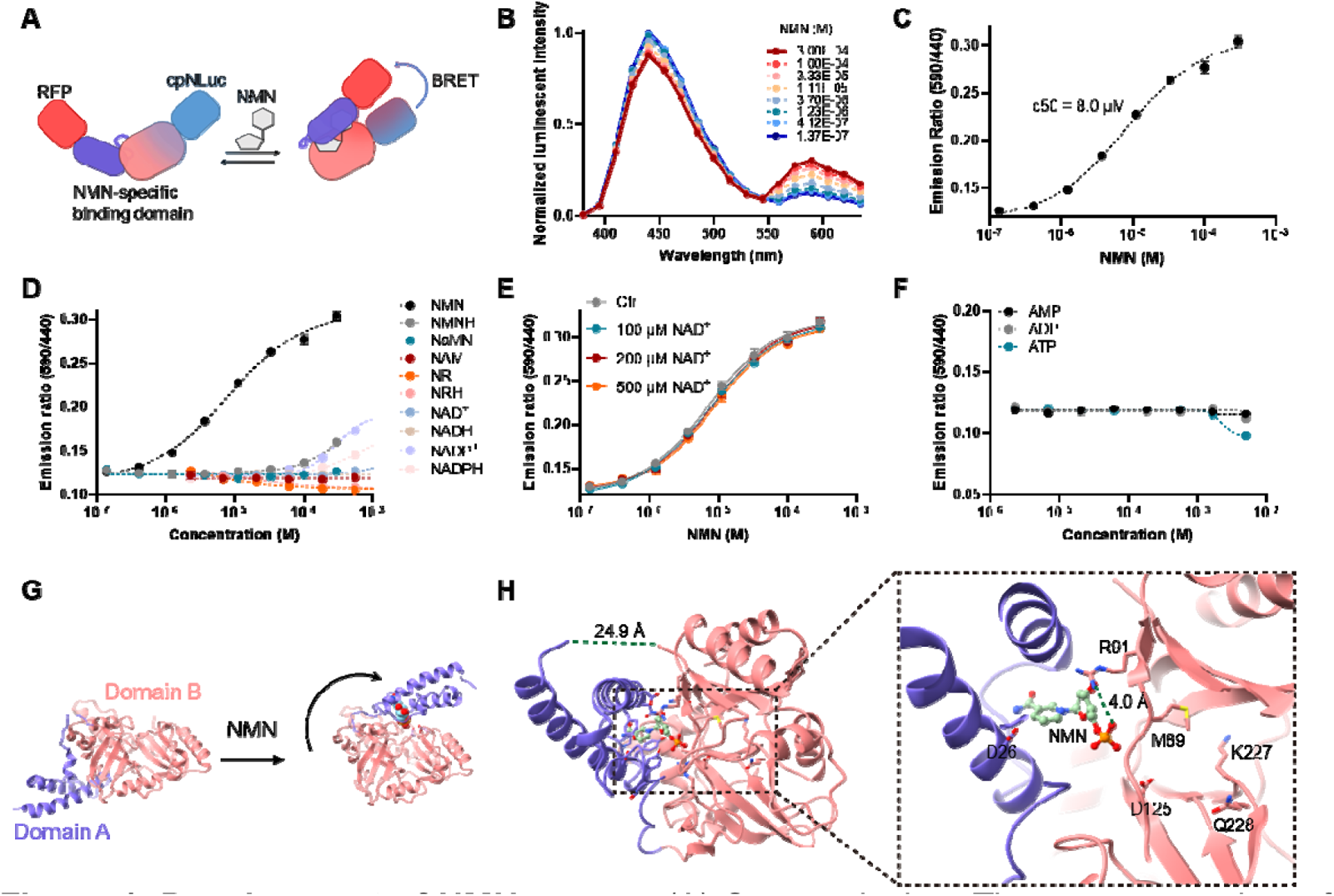
Development of NMN sensor. (A) Sensor design: The sensor consists of cpNLuc, mutated *Ef*LigA and mScarlet-I. The binding of NMN induces conformational change in *Ef*LigA and pulls cpNLuc and mScarlet-I in proximity to enable BRET. (B) Emission spectra of NMoRI at various concentrations of NMN. (C) Response curve of NMoRI towards NMN. (D) Titration of NMoRI with NMN, NMNH, NaMN, NAM, NR, NRH, NAD^+^, NADH, NADP^+^, and NADPH. (E) Titration of NMoRI with NMN in presence of various concentrations of NAD^+^. Additional NAD^+^ does not affect NMoRI’s response towards NMN. (F) Titration of NMoRI with AMP, ADP, and ATP (AXPs). NMoRI is not responsive towards AXPs. (G) Proposed conformational change of *Ef*LigA induced by NMN binding (PDB code: 1TA8). (H) 3D structure of engineered *Ef*LigA used in NMoRI (PDB code: 8JYD) with bound NMN by X-ray crystallography. Right panel details the NMN binding site and mutated amino acid residues shown in sticks. In (C), (D), (E), (F), data are shown as mean ± SD from at least three independent measurements.

To confirm that NMN binding induced conformational changes in the engineered sensor, we determined the structure of the NMoRI’s NMN-sensing domain by X-ray diffraction (Table S1). The distance between the N-terminus (L6) and C-terminus (P317) changed from 59.8 Å to 24.9 Å in the *Ef*LigA variant of NMoRI (Figure S1B, 1G-H), indicating that NMN indeed prompted a substantial conformational change in the sensor. The details of the NMN binding pocket suggested that the ligand could interact with R91 through electrostatic interactions (Figure 1H). In summary, we successfully developed a highly specific NMN sensor through rational engineering of *Ef*LigA variants.

### NMoRI records dynamics of NMN uptake in live cells

NMN uptake in mammalian cells is believed to occur through the NR-NRK pathway and/or the Slc12a8 transporter (Figure 2A).^9^ To investigate the dynamics of NMN uptake, we introduced a cytosol-specific NMoRI (Cyto-NMoRI) into HEK 293T cells and monitored the cytosolic NMN changes with presence of different NAD^+^ precursors. We found that the BRET signal of Cyto-NMoRI increased rapidly within 4 minutes in response to NMN and reached a plateau thereafter (Figure 2B). Notably, cells incubated with 100 μM NMN exhibited a faster and higher response compared to those with 10 μM NMN. We also recorded the response of Cyto-NMoRI to other precursors. As expected, nicotinamide riboside (NR) and dihydronicotinamide riboside (NRH) also caused rapid increases in the BRET ratio, suggesting that their fast uptake and conversion to NMN (Figure 2C). However, 100 μM nicotinamide (NAM) failed to increase the BRET ratio of Cyto-NMoRI within 10 minutes (Figure 2C), indicating the delayed catalytic conversion from NAM to NMN by nicotinamide phosphoribosyltransferase (NAMPT).^24^ Among the tested conditions, extracellular NMN is the most efficient precursor in boosting cytosolic NMN (Figure 2D). Furthermore, we introduced Cyto-NMoRI into 3T3-L1 and HepG2 cells and found that they responded to NMN and NR similarly to HEK 293T cells (Figure S2A-D). However, there was no significant change observed after incubating HepG2 cells with NRH (Figure S2D).

**Figure 2.**
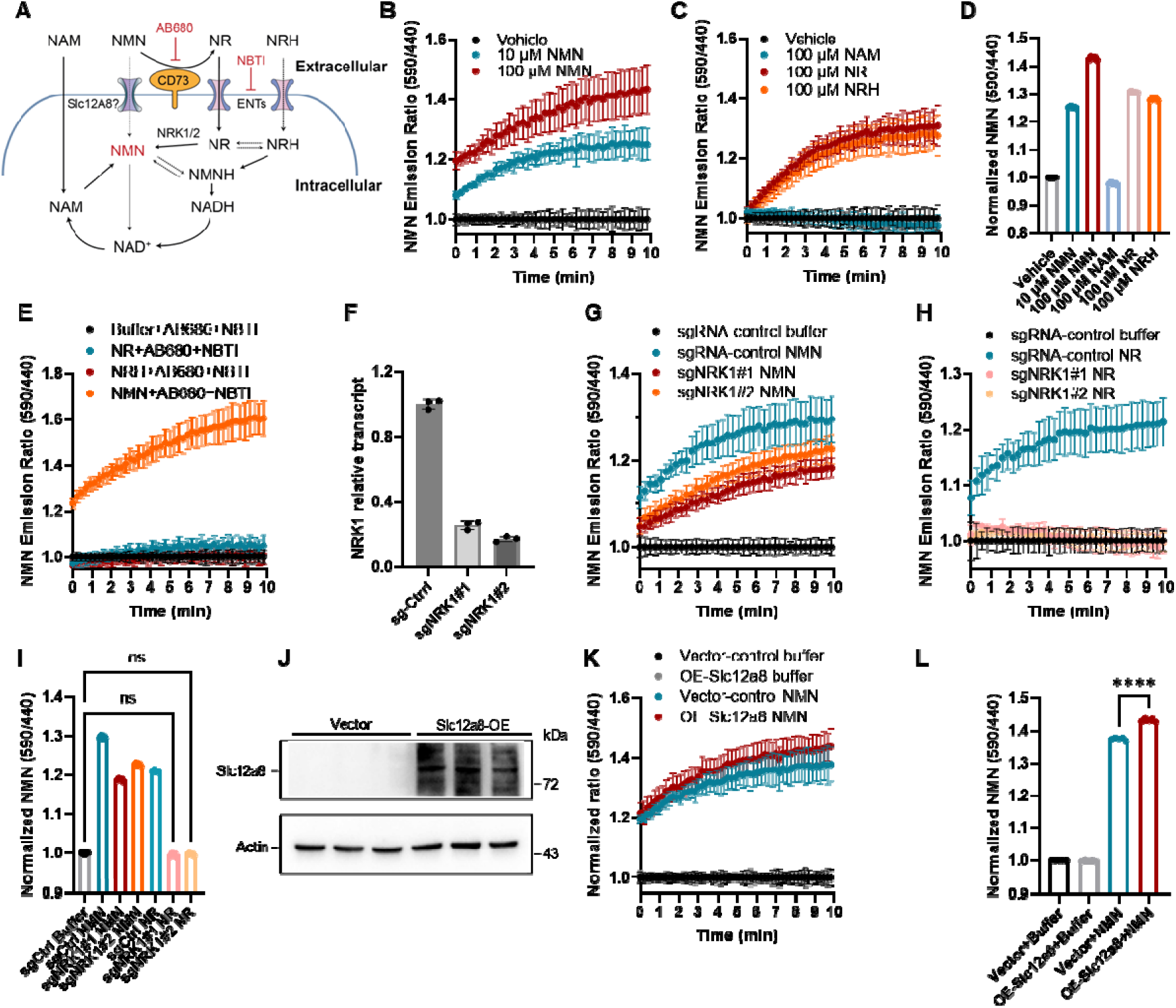
Uptake dynamics of NMN and derivatives in HEK 293T cells. (A) Major transport pathways of NAD^+^ precursors in mammalian cell. (B) Time courses of NMoRI BRET ratios from HEK 293T cells exposed to 0, 10 and 100 μM NMN. (C) Time courses of NMoRI BRET ratios from HEK 293T cells exposed to 100 μM NAM, NR, and NRH. (D) Normalized endpoint NMoRI BRET ratios induced by NAD^+^ precursors. NMN, NR and NRH effectively increased NMN levels in HEK293T cells. (E) Time courses of NMoRI BRET ratios from HEK 293T cells treated with an inhibitor cocktail of 5 μM AB680 and 10 μM NBTI for 1 h and then incubated with 100 μM NMN, or 100 μM NR, or 100 μM NRH. (F) Relative transcript levels of HEK 293T *NRK1*. (G) Time courses of NMoRI BRET ratios from HEK 293T cells incubated with 100 μM NMN after NRK1 knockdown. (H) Time courses of NMoRI BRET ratios from HEK 293T cells incubated with 100 μM NR after NRK1 knockdown. (I) Normalized endpoint NMoRI BRET ratios after treatment according to figure (G) and (H). (J) Western blot of Slc12a8 in HEK 293T after transfection by empty vector or Slc12a8 overexpression vector. (K) Time courses of NMoRI BRET ratios from HEK 293T cells with Slc12a8 overexpression incubated with 100 μM NMN. (L) Normalized endpoint NMoRI BRET ratios after treatment according to figure (K). Data are shown as mean ± SD from at least six independent measurements. Statistical significance was calculated by one-way ANOVA analysis followed by Dunnett’s multiple comparison in (I) or unpaired t test in (L). ns, *P* > 0.05; *, *P* < 0.05; **, *P* < 0.01; ***, *P* < 0.001; ****, *P* < 0.0001.

To investigate the NR-NRK conversion mechanism of NMN in HEK 293T cells, we used AB680 and NBTI to inhibit CD73 and ENTs (Figure 2A) with the presence of 100 μM NR or NMN. These inhibitors significantly suppressed the response of HEK 293T cells towards NR, resulting in only a slight increase of cellular NMN after 4 minutes (Figure 2E). However, the NMoRI’s signal increased rapidly with NMN despite of CD73 and ENTs inhibition, indicating NR-independent pathways for cellular NMN uptake. In addition, we knocked down *NRK1* (Figure 2F), the primary kinase responsible for intracellular conversion from NR to NMN.^25^ The increase of cytosolic NMN in HEK 293T cells incubated with extracellular NMN was slightly inhibited in *NRK1* knockdown cells (Figure 2G), whereas the response to NR was completely abolished (Figure 2H, I), further confirming the multiple mechanisms of NMN uptake. To investigate the role of Slc12a8 in NMN transportation, we overexpressed *Slc12a8* gene in HEK 293T cells containing Cyto-NMoRI (Figure 2J). The Slc12a8 overexpression led to an increased rate of cytosolic NMN concentration change after adding 100 μM of extracellular NMN (Figure 2K, L).

Taking together, we have captured for the first time the rapid modulatory dynamics of cytosolic NMN by NAD^+^ precursors where we showed that NMN uptake in HEK 293T cells is not solely dependent on NR-NRK conversion and that NR and NMN employ distinct mechanisms for NAD^+^ regulation.

### NMoRI reports distinct subcellular NMN signatures of NAD^+^ precursors

Enhancing cellular and tissue NAD^+^ by NAD^+^ precursors is well-established and believed to be beneficial for a range of age-related diseases. Given that NMN is a primary precursor of NAD^+^ (Figure 3A), mapping subcellular NMN is important for the comprehensive understanding of NAD^+^ metabolism. We hence developed HEK 293T cells that stably express NMoRI in cytosol (Cyto-NMoRI), nuclear (Nuc-NMoRI), and mitochondria (Mito-NMoRI) by adding organelle-targeting peptides to the N-terminus of the sensor (Figure S3). Currently, the subcellular free NMN levels remain uncharacterized. To evaluate the NMoRI’s utility for quantifying intracellular free NMN, we established calibration curves for HEK 293T cells expressing the organelle-specific sensors (Figure 3B-D). The sensors reported free NMN levels at 3.07 μM in cytosol, 3.65 μM in nucleus, and less than 0.1 μM in mitochondria. These values were in line with previously reported measurements using LC-MS.^26^ Notably, The very low mitochondrial NMN indicated a distinct environment for NMN-NAD^+^ metabolism in the organelle.

**Figure 3.**
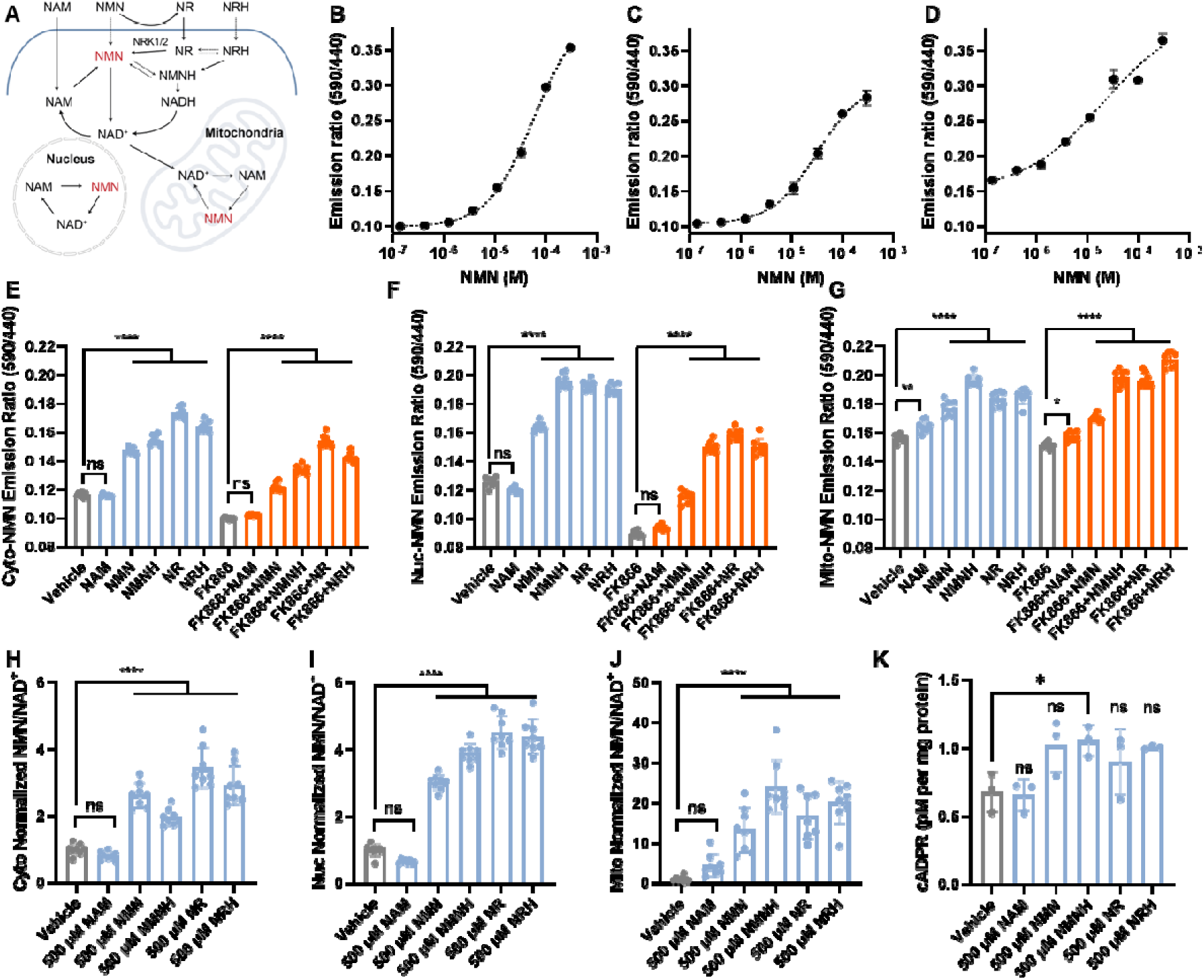
NMoRI detects NMN and NMN/NAD^+^ ratios in subcellular compartments. (A) Schematic of subcellular NMN and NAD^+^ metabolism. Calibration curves for HEK 293T cells stably expressing NMoRI in cytosol (B), nucleus (C), and mitochondria (D), n ≥ 6. Effects of 500 μM NAD^+^ precursors on cytosolic (E), nuclear (F), and mitochondrial (G) NMN levels with and without FK866 (10 nM), n ≥ 6. Effects of 500 μM NAD^+^ precursors on cytosolic (H), nuclear (I), and mitochondrial (J) NMN/NAD^+^ levels, n ≥ 6. (K) Quantification of cADPR (pM per mg protein) produced from HEK 293T cells treated with 500 μM NAD^+^ precursors for 6h. Data are shown as mean ± SD. Statistical analysis was performed by one-way ANOVA analysis followed by Dunnett’s multiple comparison. ns, *P* > 0.05; *, *P* < 0.05; **, *P* < 0.01; ***, *P* < 0.001; ****, *P* < 0.0001.

Furthermore, we showed that NAD^+^ precursors, including NMN, dihydronicotinamide mononucleotide (NMNH), NR, and NRH, can effectively augment NMN levels in all tested organelles after 6 hours of incubation (Figure 3E-G). Notably, NR exhibited the highest efficacy in boosting NMN in cytosol (Figure 3E) after 6 hours, while NMNH is the most effective in nuclear (Figure 3F) and mitochondria (Figure 3G). Surprisingly, NAM did not affect NMN in neither cytosol (Figure 3E) nor nucleus (Figure 3F) but increased mitochondrial NMN (Figure 3G). In NAD^+^-depleted cells (treated by FK866), the boosting effects of NMN, NMNH, NR, and NRH remained potent for all organelles (Figure 3E-G), while NAM was effective only in mitochondria (Figure 3G).

Recent studies highlighted the pivotal role of increased NMN/NAD^+^ ratio in activating SARM1,^8^ a key regulator for neurological diseases.^27^ NAD^+^ precursors such as NR and NMN hold the promise of improving age-related diseases via NAD^+^ pathways. Other novel precursors such as NRH and NMNH claim to be highly potent in boosting NAD^+^. But much is unknown for their effects on disturbing intracellular NMN/NAD^+^ ratios, leaving the potential risk of SARM1 activation uncharacterized. We hance quantified the NMN/NAD^+^ ratio in different subcellular compartments after supplementing these precursors, with organelle-specific NAD^+^ levels measured by the previously developed NAD^+^ sensor NS-Olive (Figure S4). The administration of NAM did not significantly alter the NMN/NAD^+^ values in all tested subcellular compartments (Figure 3H-J). Meanwhile, NMN, NMNH, NR and NRH significantly increased the NMN/NAD^+^ ratios, with the most pronounced effect observed in mitochondria. We further evaluated the consequences of altered NMN/NAD^+^ ratios by testing the intracellular cADPR levels as indicator for SARM1 activation.^28,29^ For NMNH treated HEK293T cells, cADPR is slightly increased (Figure 3K). Other precursors also induced slight cADPR increase but with no significance. It is worth noticing that HEK293T cells have intrinsically low SARM1 expression levels. The high NMN/NAD^+^ ratio induced activation could be more pronounced for other cells with high SARM1 levels.^29^

In summary, NMoRI quantified the organelle specific NMN and NMN/NAD^+^ signature of various NAD^+^ precursors. Given the naturally low mitochondrial NMN levels, many NAD^+^ precursors significantly expand the mitochondrial NMN/NAD^+^ ratio by 10 to 30 folds, leading to SARM1 activation for certain precursors. Hence, the live cell NMN/NAD^+^ ratio measured by NMoRI and NS-Olive provided an important perspective to comprehensively evaluate the biological consequences of NAD^+^ modulating strategies.

### Quantifying modulations of subcellular NMN/NAD^+^ by key NAD^+^ enzymes

Nicotinamide mononucleotide adenylyltransferases (NMNATs) and solute carrier family 25 member 51 (Slc25a51) are important proteins for subcellular NAD^+^ and NMN homeostasis, with NMNAT1 localized in nucleus, NMANT2 in cytosol, NMANT3 in mitochondria, and Slc25a51 for mitochondrial NAD^+^ transport (Figure 4A). These proteins have been reported to regulate processes such as ADP-ribosylation and mitochondrial NAD^+^ transportation. We hence investigated the levels of subcellular NMN, NAD^+^ and NMN/NAD^+^ ratios after their knockdown and overexpression of in cytosol, nuclear and mitochondria.

**Figure 4.**
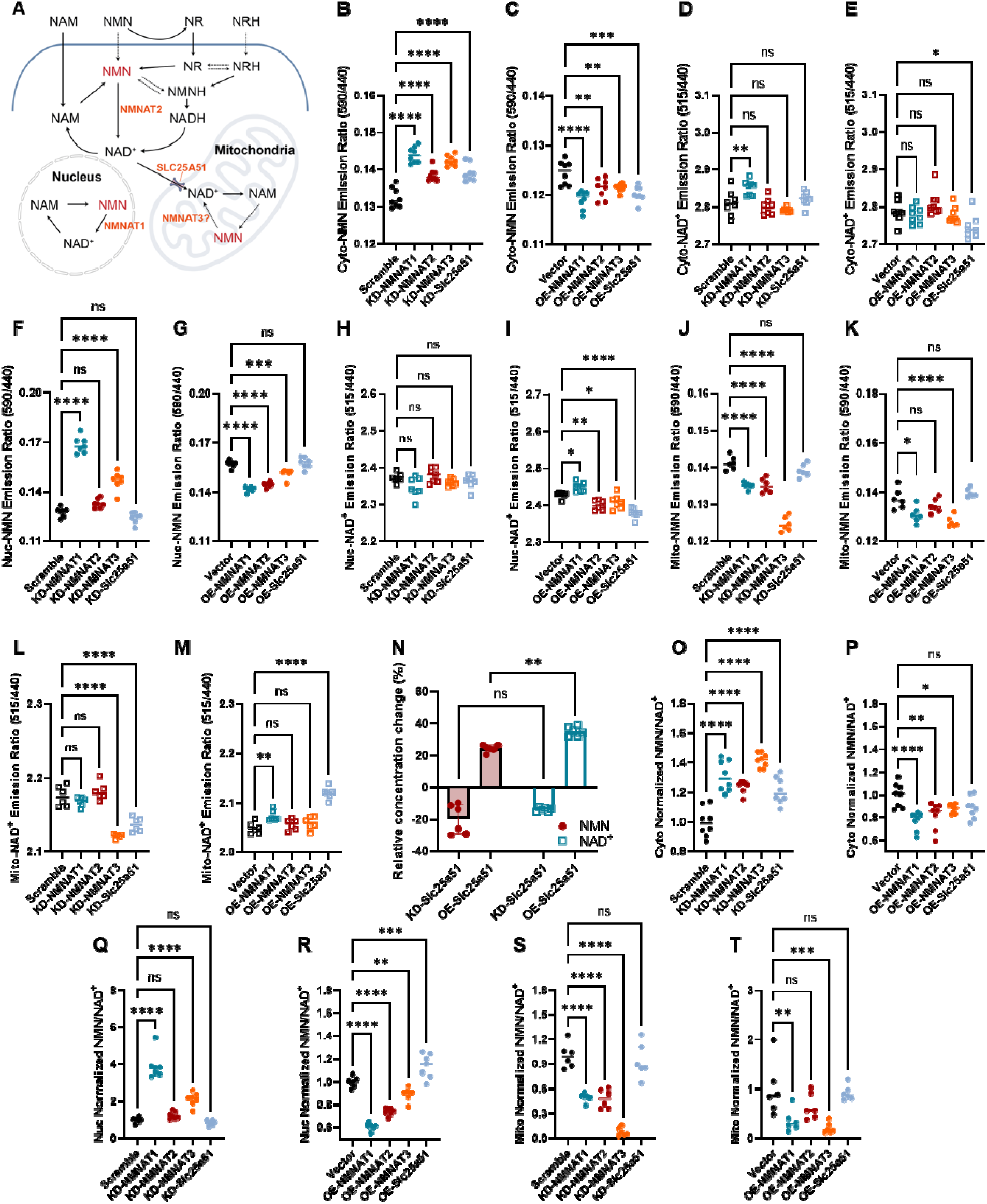
Subcellular NMN regulation in HEK 293T cells. (A) Schematic representation of regulatory network of NMN metabolism. Effects of knockdown (B) and overexpression (C) of selected genes on cytosolic NMN BRET ratio, n ≥ 6. Effects of knockdown (D) and overexpression (E) of selected genes on cytosolic NAD^+^ BRET ratio, n ≥ 6. Effects of knockdown (F) and overexpression (G) of selected genes on nuclear NMN BRET ratio, n ≥ 6. Effects of knockdown (H) and overexpression (I) of selected genes on nuclear NAD^+^ BRET ratio, n ≥ 6. Effects of knockdown (J) and overexpression (K) of selected genes on mitochondrial NMN BRET ratio, n ≥ 6. Effects of knockdown (L) and overexpression (M) of selected genes on mitochondrial NAD^+^ BRET ratio, n ≥ 6. (N) Relative concentration changes of mitochondrial NMN and NAD^+^ in HEK 293T after Slc25a51 knockdown or overexpression compared to scramble RNA or empty vector. Effects of knockdown (O) and overexpression (P) of selected genes on cytosolic NMN/NAD^+^ ratio, n ≥ 6. Effects of knockdown (Q) or overexpression (R) of selected genes on nuclear NMN/NAD^+^ ratio, n ≥ 6. Effects of knockdown (S) or overexpression (T) of selected genes on mitochondrial NMN/NAD^+^ ratio, n ≥ 6. Error bars represent SD. Statistical analysis was performed by one-way ANOVA analysis followed by Dunnett’s multiple comparison (B-M and O-T) or two-way ANOVA analysis followed by Sidak’s multiple comparison (N). ns, *P* > 0.05; *, *P* < 0.05; **, *P* < 0.01; ***, *P* < 0.001; ****, *P* < 0.0001.

In cytosol, the knockdown of NMN-converting enzymes NMNAT1-3 significantly upregulated the free cytosolic NMN levels (Figure 4B, S5), and their overexpression reduced the free NMN (Figure 4C, S5), demonstrating their direct NMN-converting nature. On the other hand, the cytosolic NAD^+^ levels were augmented by knocking down NMNAT1 but not NMNAT2 nor NMNAT3, indicating the competing roles of nuclear and cytosolic NMNATs (Figure 4D, S6). Moreover, Slc25a51 affected both NMN and NAD^+^ pools in the cytosol with its knockdown upregulates cytosolic NMN (Figure 4B) and its overexpression reduces cytosolic NMN (Figure 4C) and NAD^+^ (Figure 4E).

Similarly in nucleus, knocking down NMNAT1 and NMNAT3, but not NMNAT2, significantly increased nuclear NMN levels (Figure 4F), while overexpressing NMNAT1-3 reduced the nuclear NMN with NMNAT1 showing the strongest effect (Figure 4G). Moreover, knocking down NMNAT1-3 did not affect nuclear NAD^+^ (Figure 4H). But the overexpression of NMNAT1 increased NAD^+^, and NMNAT2 and 3 reduced NAD^+^ in the nucleus (Figure 4I), underlining the competition between NMNATs in different cell compartments. Slc25a51 did not show much effect on nuclear NMN (Figure 4F, G), but its overexpression significantly reduced nuclear NAD^+^ (Figure 4I).

For mitochondria, NMNAT3, among all tested NMNATs, exhibited the strongest impact on mitochondrial NMN (Figure 4J, K) and NAD^+^ (Figure 4L). Interestingly, the nuclear-localized NMNAT1 showed measurable effects on mitochondrial NMN and NAD^+^ where the cytosolic NMNAT2 did not (Figure 4K, M), indicating a potentially more active interaction between the nuclear and mitochondrial NMN and NAD^+^ metabolism. Slc25a51 significantly affected only the mitochondrial NAD^+^ (Figure 4L, M) but not NMN (Figure 4J, K). Moreover, Slc25a51 affected more mitochondrial NAD^+^ than NMN in terms of concentration changes (Figure 4N), indicating its role in mitochondrial NAD^+^ transportation.

We further quantified the effect of these genes on NMN/NAD^+^ ratios. As expected, knocking down NMNAT1-3 significantly increased cytosolic NMN/NAD^+^ ratios (Figure 4O) and their overexpression reduced such ratio (Figure 4P). NMNAT1 is the most influential enzyme on NMN/NAD^+^ ratios in nuclear (Figure 4Q, R), and NMNAT3 in mitochondria (Figure 4S, T). Modulation of Slc25a51 showed non-significant and dispersed effects on NMN/NAD^+^ ratios in mitochondria (Figure 4S, T), but affected cytosolic and nuclear NMN/NAD^+^ ratios (Figure 4O, R).

Here, we quantified the effects of NMNAT1-3 and Slc25a51 on NMN, NAD^+^ and NMN/NAD^+^ ratios in live cell compartments using NMoRI. The effect of NMNAT1-3 was in line with their NMN converting activities and localization. Interestingly, the unexpected effects of mitochondrial and nuclear NMNATs on each other’s NMN and NAD^+^ pools indicated potentially active material exchanges between mitochondria and nuclear. Furthermore, we showed that mitochondrial NAD^+^ transporter Slc25a51 significantly affected mitochondrial NAD^+^ but not NMN, nor NMN/NAD^+^ ratios. Collectively, we demonstrated the ability of NMoRI in indicating organelle specific NMN metabolism and in dissecting functions of key enzymes for NMN and NAD^+^ homeostasis.

### NMN/NAD^+^ ratio indicates NMN transport between cytosol and mitochondria

An important question in NAD^+^ metabolism is whether there exist NMN transporters in mitochondria (Figure 5A). Here, we sought to identify potential NMN transporters by monitoring the mitochondrial NMN, NAD^+^ and NMN/NAD^+^ ratio after genetically modulating speculated solute carrier proteins.

**Figure 5.**
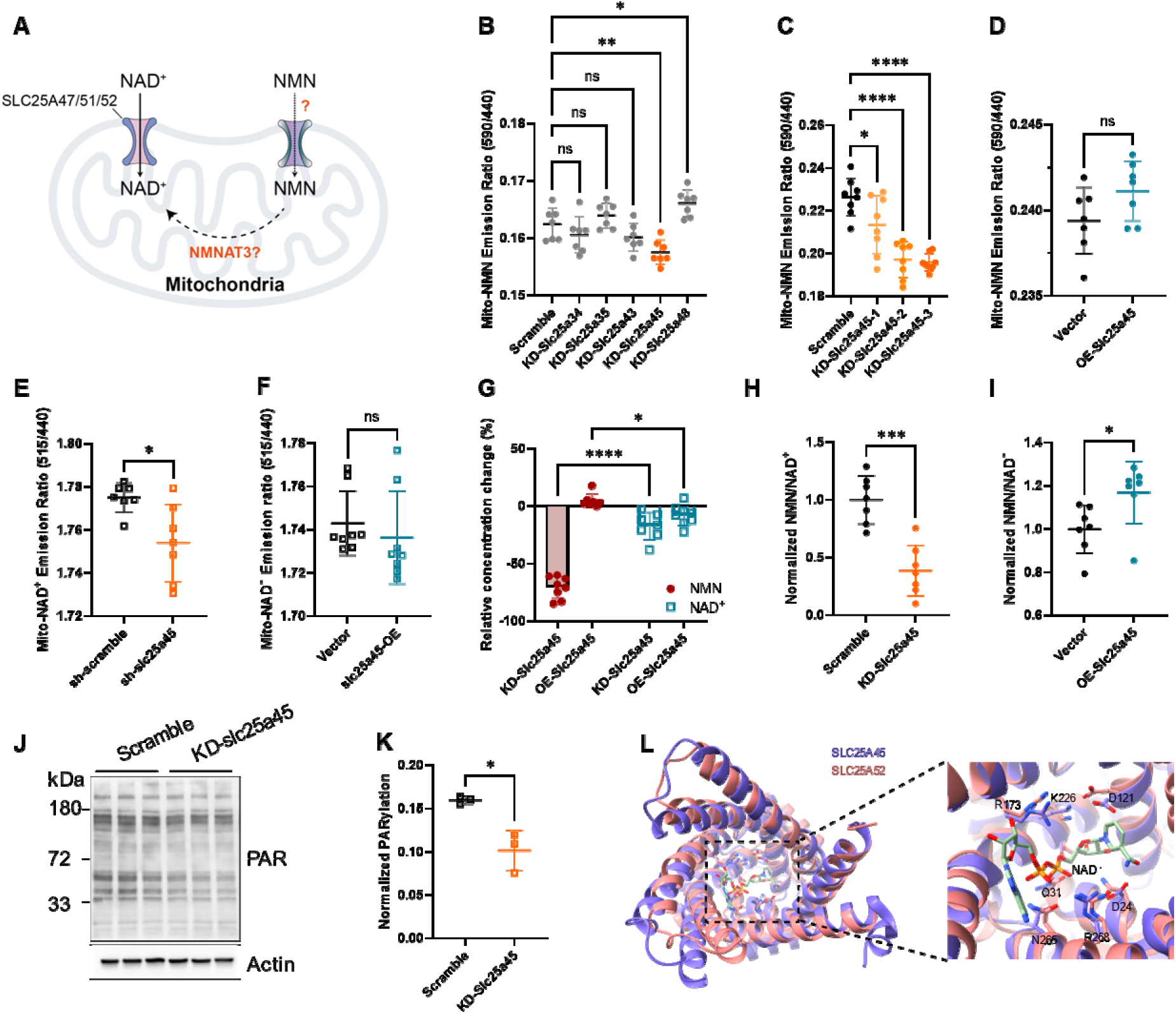
Discovering potential mitochondrial NMN transporter. (A) Schematic representation of NMN metabolism in mitochondria. (B) Screening for modulators of mitochondrial NMN by knocking down mitochondrial solute carrier family proteins in HEK 293T cells. NMN was monitored by Mito-NMoRI. (C) Effects of knocking down Slc25a45 on mitochondrial NMN in HEK 293T cells by shRNA KD-Slc25a45#1, #2, and #3. (D) Effects of overexpressing Slc25a45 on mitochondrial NMN in HEK 293T cells. (E) Effects of knocking down Slc25a45 on mitochondrial NAD^+^ in HEK 293T cells, KD-Slc25a45 denote KD-Slc25a45#3 here and follows. (F) Effects of overexpressing Slc25a45 on mitochondrial NAD^+^ in HEK 293T cells. (G) Relative concentration changes of mitochondrial NMN and NAD^+^ in HEK 293T after Slc25a45 knockdown or overexpression. (H) Effects of knocking down Slc25a45 on mitochondrial NMN/NAD^+^ ratio in HEK 293T cells. (I) Effects of overexpressing Slc25a45 on mitochondrial NMN/NAD^+^ ratio. (J) PAR immunoblot analysis of lysate from HEK 293T cells stably expressing mitochondria targeted PARP1 cd sensor. PARylation levels reflect NAD^+^ availability. (K) Statistics of normalized PARylation level in mitochondria with and without Slc25a45 knockdown. (L) Binding pocket of Slc25a45 and the mitochondrial NAD^+^ transporter Slc25a52 models. Error bars represents SD. Statistical analysis was performed by one-way ANOVA analysis followed by Dunnett’s multiple comparison (B, C), two-way ANOVA analysis followed by Sidak’s multiple comparison (G) or unpaired t test (D-F, H-I, and K). ns, *P* > 0.05; *, *P* < 0.05; **, *P* < 0.01; ***, *P* < 0.001; ****, *P* < 0.0001.

We knocked down Slc25a34, 35, 43, 45 and 48 to screen for their effects on mitochondrial NMN indicated by Mito-NMoRI in HEK 293T cells. Notably, the knockdown of Slc25a45 significantly decreased mitochondrial NMN in both the screening and confirmational experiments (Figure 5B and 5C). The overexpression, however, did not significantly affect mitochondrial NMN (Figure 5D). To further confirm the regulatory role of Slc25a45, we measured mitochondrial NAD^+^ levels and showed that knocking down Slc25a45 moderately reduced NAD^+^ (Figure 5E), while its over expression showed no significant effect (Figure 5F). Compared to the previous identified mitochondrial NAD^+^ transporter Slc25a51 (Figure 4N), Slc25a45 showed here a much more pronounced effect on NMN than on NAD^+^ (Figure 5G), indicating its role as a potential NMN transporter. We then measured the NMN/NAD^+^ ratio change induced by Slc25a45 knockdown and overexpression. As expected, knocking down Slc25a45 decreased NMN/NAD^+^ ratio and its overexpression led to increased NMN/NAD^+^ ratio.

To further confirm the regulatory role of Slc25a45 on NAD^+^ metabolism, we employed PARPcd, another NAD^+^ indicator, to quantify Slc25a45’s effect on mitochondrial NAD^+^.^23,30^ Similarly, PARPcd assay showed that knocking down Slc25a45 decreased mitochondrial NAD^+^ (Figure 5J, K), suggesting that Slc25a45 indeed plays pivotal roles in NMN and NAD^+^ metabolism. We then performed a structural alignment between Slc25a45 and the NAD^+^ transporter Slc25a52 using models from AlphaFold protein database. The alignment revealed that key residues interacting with NAD^+^ and NMN moieties are conserved (Figure 5G). In summary, our findings unveiled mitochondrial membrane protein Slc25a45 as a potential transporter of mitochondrial NMN.

## DISCUSSION

NMN plays critical roles in NAD^+^ metabolism and has been shown to exert pharmacological activities in various pathologies.^31–33^ However, its metabolic dynamics in subcellular compartments remain uncharacterized. In this study, we successfully designed the first genetically encoded NMN sensor for its quantification in live cells. To accomplish this, we took protein domains from *Ef*LigA as the analyte binding scaffold and combined rational design and saturated mutagenesis to engineer its specificity towards NMN. As a result, we developed BRET NMN sensors named NMoRIs with c50s ranging from 1 - 30 μM, making them suitable for various quantification scenarios. Using NMoRI, we explored the dynamics of different mechanisms for NMN uptake. We then quantified subcellular NMN and precursors’ perturbations on NMN/NAD^+^ ratios; and identified a potential mitochondrial NMN transporter.

Whether cells directly transport NMN remains an important question for understanding NAD^+^ metabolism. Previous studies suggested that NMN enters cells via an indirect pathway, involving its conversion first into NR by CD73, followed by NR’s transport into cells through nucleoside transporters, and a final reconversion into NMN by NR kinase.^25,34–38^ However, some studies indicated that NMN is poorly catalyzed by CD73,^39^ and that Slc12a8 functions as a NMN transporter.^40,41^ Due to the lack of live cell quantification methods for NMN, details of its cellular uptake remain obscure. Our NMN sensors provided the first opportunity to detect cellular NMN in real-time. We observed that NMN can induce immediate and dose-dependent cellular responses. NR and NRH also induced significant NMN increase in HEK 293T cells within minutes despite of a slower rate, suggesting that NR and NRH can be transported into HEK 293T cells and rapidly converted into NMN. Pharmacological inhibition of CD73 and ENTs, as well as genetic knockdown of NRK1, completely abolished NR uptake and conversion into NMN by HEK 293T cells but did not stop the uptake of extracellular NMN. Moreover, overexpression Slc12a8 slightly increased NMN uptake in HEK 293T cells. Based on these observations, we speculate that NMN can be directly taken up by HEK 293T cells, meanwhile the indirect NR-NRK pathway also contributes to the NMN uptake. However, we also observed cell-type specific responses towards the different precursors, suggesting different mechanisms could be emphasized in cell-specific manners. In future, NMoRI would facilitate further investigations into NMN uptake mechanisms in different cells, tissues, and pathological events.

Organelle specific NMN concentrations in live cells were not available before.^9^ Based on the established *in cellulo* titration curves, we observed that cytosol and nucleus maintained a basal NMN level between 3 μM to 4 μM, while the free NMN level in mitochondria was less than 0.1 μM. This is in agreement with previous reported values obtained from cell lysates in other cell lines.^26^ The NMN/NAD^+^ ratio is known to be critical for SARM1 activation, which regulates axon degeneration. Given the increasing interest in boosting NAD^+^ through precursors to counter age-related diseases, it is important to assess their impacts on cellular NMN/NAD^+^ ratios. NMoRI now makes it possible in combination with previously designed NAD^+^ sensors. In HEK 293T cells, we found that NMN, NMNH, NR, and NRH can simultaneously increase NMN, NAD^+^ and NMN/NAD^+^ ratios in all tested subcellular compartments. Remarkably, the mitochondrial NMN/NAD^+^ ratio can increase by more than 20 folds, potentially activating SARM1.^8^ However, further research is required to confirm the downstream biological effects from elevated NMN/NAD^+^ ratios.

Mitochondria is unique for NAD^+^ and NMN metabolism because of its very low NMN levels and distinct regulatory mechanisms. Recent studies have identified a mitochondrial NAD^+^ transporter, but how NMN enters mitochondria remains unclear. The mitochondrial localization of NMNAT3 strongly suggests that mitochondria may take up NMN. Yet no study has confirmed NAD^+^ being converted from NMN in isolated mitochondria.^42,43^ The discovery of mitochondrial NAD^+^ transporters further supports the direct transport of mitochondrial NAD^+^ from cytosol. However, the knockdown of NMNAT3 also decreases mitochondrial NAD^+^,^22^ implying that NMN contributes to the mitochondrial NAD^+^ pool. Using NMoRI, we showed that knocking down mitochondrial membrane protein Slc25a45 considerably reduced mitochondrial NMN and affected NMN/NAD^+^ ratio, exerting much more pronounced effect on NMN than on NAD^+^. By contrast, the identified NAD^+^ transporter only affect mitochondrial NAD^+^ but not NMN. Overall, the observation indicates that Slc25a45 is a potential NMN transporter.

In summary, we engineered an NMN-specific bioluminescent sensor protein to quantify subcellular NMN dynamics and regulations in live cells. As exemplified in this work, the sensor should lead to the exploration of many important questions in NMN and NAD^+^ metabolism.

## Methods

### Molecular biology

A pET51b vector was used for sensor protein production in *E. coli* BL21 (DE3). A pCDH-CMV-MCS-EF1-Neo vector or pLV3-CMV-MCS-Puro vector were used for generating sensor-expressing HEK 293T cells. The initial sensor-encoding plasmid was constructed as previous described.^23^ The genes encoding NMNAT1, NMNAT2, NMNAT3, Slc25a51, and Slc12a8 were synthesized by TSINKE Biotech and subcloned into pLV3-CMV-MCS-Puro vector with a 16-nt overlap for seamless cloning. The sensor genes were amplificated with 16-nt overlap and cloned into pCDH-CMV-MCS-EF1-Neo vector using seamless cloning. The cytosol-targeting sequence (NES), mitochondrial targeting sequence (2 x COXVIII), and nuclear targeting sequence (NTS) were added to the N terminus of the sensor for specific expression in cytosol, mitochondria, and nucleus.

NES: ATGCTGCAGAATGAACTGGCACTGAAGCTCGCAGGCCTGGACATTAACAAGACCGGTGGAAGCGGC 2 x COXVIII: ATGTCTGTTCTGACTCCTCTGCTGCTCCGGGGTCTCACAGGTTCCGCAAGAAGACTCCCCGTGCCTAGGGCCAAAATTCATTCACTGGGGGACCCCATGAGCGTGCTCACCCCACTCCTGCTGCGGGGGCTGACCGGCAGCGCTAGGCGGCTGCCAGTCCCCAGGGCCAAGATCCACAGTCTCGGCGATCCCAAG NTS: ATGGATCCAAAAAAGAAGAGAAAGGTA

Sensor variants were generated by site-directed mutagenesis with specific oligonucleotide primers purchased from TSINKE Biotech. Amplification was performed using KOD plus neo polymerase (Toyobo). The sequences and resulting vector maps of the mutagenesis library are available upon request. Each variant was confirmed by sequencing.

### Protein expression and purification

All sensor variants and the *Ef*LigA domain variants were expressed in *E. coli* BL21 (DE3). The *E. coli* was cultured in 200 mL LB medium containing 50 μg/mL ampicillin at 37°C until the cultures reached an OD of 0.6. Then, the growth temperature was shifted to 25°C and protein expression was induced by adding 0.2 mM IPTG. After 16 h of incubation, bacteria were harvested and suspended in 50 mM Tris-HCl buffer, pH 8.0, containing 500 mM NaCl and 10 mM imidazole, and lysed via high pressure homogenizer. Cell lysate was centrifuged at 12,000 × *g* for 15 min at 4°C, and the supernatant was loaded sequentially onto Ni-NTA (Smart Lifesciences Inc., China) and Strep-Tactin columns (Smart Lifesciences Inc., China) for purification. The purified protein was desalted and exchanged into 50 mM HEPES buffer containing 50 mM NaCl (pH 7.2) with an Amicon Ultra-15 centrifugal filter (10 kDa MWCO, Merk Millipore Inc.). Protein concentration was determined through BCA assay and subsequently diluted to the requisite concentration for subsequent *in vitro* assays.

### Protein crystallization and structure determination

For protein crystallization, the *Ef*LigA domain of NMoRI was further purified by gel filtration using a Superdex-200 column (GE Healthcare) equilibrated against buffer (25 mM Tris-HCl pH 7.4, 100 mM NaCl). The pure proteins were concentrated with an Amicon Ultra-15 centrifugal filter (10 kDa MWCO, Merk Millipore Inc.) to around 20 mg/mL.

Crystallization experiments were conducted in 48-well plates by the hanging-drop vapor diffusion method at 293 K, and each hanging drop was prepared by mixing 1.0 μL each of protein solution (containing 10 mM NMN) and reservoir solution. The reservoir solution used for successful crystal growth contained 0.1 M MES pH 6.0 and 20% PEG mono 2000. Crystals were cryoprotected using a reservoir solution containing 25% (v/v) ethylene glycol. The X-ray data were collected on 19U beamline at the Shanghai Synchrotron Radiation Facility. Data were indexed, integrated and scaled using xia2-3dii.^44^ The structure was solved by molecular replacement method using the crystal structure of wild type *Ef*LigA (PDB code: 1TAE) as template. Subsequently, the structure was refined by using REFMAC5^45^ program and manual inspected with COOT.^46^ Data collection, processing, and refinement statistics were summarized in Table S1. Graphic representations of structures were generated in ChimeraX.^47^ Atomic coordinate and structure factors have been deposited in the PDB under accession code 8JYD.

### Sensor titration

The sensors were titrated with NMN, NMNH, NaMN, NAM, NAD^+^, NADH, NADP^+^, NADPH, NR, and NRH to assess their sensitivity and selectivity. For BRET sensors, various concentrations of the compounds were mixed with 1 - 2 nM sensor and 1,000-fold diluted furimazine (Nano-Glo Luciferase Assay Substrate, Promega) in 100 μL of buffer (50 mM HEPES, 50 mM NaCl, 1 mg/mL BSA, pH 7.2) in a white 96-well plate (Grener). Bioluminescence was measured using a FlexStation 3 Multi-Mode microplate reader or Tecan Spark multimode plate reader with NLuc emission measured at 440 nm and mScarlet-I emission at 590 nm. The ratios between NLuc and mScarlet-I emission (mScarlet-I/NLuc) were plotted against the compound concentrations. The sensor’s maximum ratio (Rmax), minimum ratio (Rmin), c50 and Hill coefficient (h) were obtained by fitting the data to the Hill-Langmuir equation.

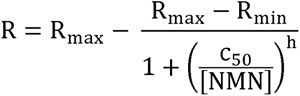

NAD^+^ concentrations in unknown samples were calculated with the rearranged the equation using the measured R.

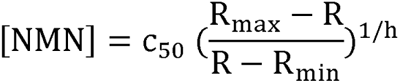

### Cell culture, lentivirus preparation and infection

HEK 293T, Hep G2, and 3T3-L1 cells were cultured in DMEM (11885, Gibco) supplemented with 10% FBS (VivaCell C04001) at 37°C with 5% CO_2_. To produce lentivirus, plasmids pMD2G, psPAX2 and the target gene plasmids (NMoRI in pCDH-CMV-MCS-EF1-Neo vector, NMNATs, Slc25a51, and Slc12a8 in pLV3-CMV-MCS-Puro vector) were employed. A 3x flag was added into the N-terminus of Slc12a8. The viral supernatant was initially filtered using a 0.45 μm PVDF filter and subsequently concentrated through PEG-8000 precipitation. The resulting viral solution was then divided into aliquots, flash frozen, and stored at -80°C util use. For cell infection, HEK 293T, HepG2, and 3T3-L1 cells cultured in 60 mm plates were exposed to the virus solutions. Subsequently, the culture media were replaced 6 h after infection. After 48 h incubation period post-infection, the cells were harvested and subjected to serial dilution to select for infected cells, employing the fluorescence of the sensors or puromycin of vector as selection marker.

### shRNA and CRISPR-Cas9-mediated gene knockdown

To create shRNAs for gene knockdown, they were ligated into a pLKO.1 puro vector. HEK 293T cells were transfected with this plasmid to generate lentivirus. Subsequently, NMoRI expressing cells were transfected with this lentivirus to induce the knockdown of targeted genes. The specific shRNA sequences used for knockdown are as follows:

1. NMNAT1 shRNA: TRCN0000035471: GTGGAAGTTGATACATGGGAA
2. NMNAT2 shRNA: TRCN0000035440: GCCAGGGATTATCTGCACAAA
3. NMNAT3 shRNA: TRCN0000035400: CAGGAAATAGTGGAGAAGTTT
4. Slc25a51 shRNA: TRCN0000060235: GCAACTTATGAGTTCTTGTTA
5. Slc25a45 shRNAs:

sh-Slc25a45#1: CCAGGTTGAAAGCTTGCAAAT

sh-Slc25a45#2: CCTCAGCTACGAATATCTCCT

sh-Slc25a45#3: GTGGTCAACTCTGTCCTGTTT

NRK1 were knocked down by using the following sgRNA target sequences: sgNRK1#1: ACATGGCCATCAAAGTATCC; sgNRK1#2: GGAAAGCGCAAGACACTCTG. The sgRNA were ligated into LentiCRISPR v2 vector. HEK 293T cells were transfected with this plasmid to generate lentivirus. Subsequently, NMoRI expressing cells were transfected with this lentivirus to induce the knockdown of targeted genes.

### Western blot

Cells were washed three times with ice-cold PBS and lysed in RIPA buffer with protease and phosphatase inhibitors. Protein samples were electrophoresed on 4-20% precast gel and transferred to a 0.45 μm PVDF membrane. Membranes were blocked with 5% no-fat milk in Tris-buffered saline containing 0.1% (v/v) Tween-20 (TBST). Primary antibodies were diluted in 1% no-fat milk in TBST and incubated at 4°C overnight. After three washes, the secondary antibody, diluted in TBST, was incubated for 1 h at RT. After another three washes, target proteins were visualized with iBright imaging system. The following primary and secondary antibodies were used for WB analysis: NMNA1 polyclonal antibody (Proteintech, 11399-1-AP), NMNAT2 polyclonal antibody (Invitrogen, PA5-115662), NMNAT3 antibody (Proteintech, 13236-1-AP), Slc25a51 antibody (ABclonal, A21105), monoclonal anti-flag M2 antibody (Millipore, F1804), anti-Actin (ABclonal, AC026), anti-Rabbit IgG (Biosharp, BL003A), anti-Mouse IgG (Biosharp, BL021A).

### RNA isolation and real-time quantitative PCR

RNA-Quick purification kit (YiShan Biotech) was used to extract total RNA from cells. Then, 500 ng total RNA was reversed transcribed into cDNA using the cDNA synthesis kit (Takara). The cDNA was diluted for suitable fold and mixed with qPCR SYBR Green Master Mix (Takara) and primer pairs. The real-time quantitative PCR was performed on 7500 real-time PCR system (Applied Biosystems). The primers used for qPCR: NRK1 Fwd: CCAAATTGCAGTGTCATATCTCAG, Rev: CCAGCAGGAAATGGCTGAC ATC. Slc25a45 Fwd: ACCCGTTTGACACTGTAAAGGTG, Rev: CAGGACAGAGTTG ACCACAGCT. Actin Fwd: CACCATTGGCAATGAGCGGTTC, Rev: AGGTCTTTGCG GATGTCCACGT.

### Cellular BRET measurement using microplate reader

Cells that stably expresse the BRET sensor were seeded at a density of 10,000 cells per well in a white 96-well plate (SPL) with DMEM culture supplemented with 10% FBS. After 24 h of incubation at 37°C with 5% CO_2_, different compounds were added into the culture medium. After 6 h of treatment, culture medium was removed by aspiration and replaced with 100 μL phenol red-free DMEM culture containing 1,000-fold diluted furimazine. Bioluminescence was measured using a FlexStation^®^ 3 Multi-Mode Microplate Reader (Molecular Devices) with NLuc emission measured at 440 nm and mScarlet-I emission at 590 nm.

### NMN, NR, and NRH uptake measurement

Cells that stably express the BRET sensor were seeded at a density of 15,000 cells per well in a white 96-well plate (SPL) with DMEM culture supplemented with 10% FBS. After 24 h of incubation at 37°C with 5% CO_2_, the inhibitors AB680 and NBTI were added to achieve final concentrations of 5 μM and 10 μM, respectively. Following an 1 h incubation, the cells were washed three times with HBSS buffer in the presence of 5 μM AB680 and 10 μM NBTI. Subsequently, the BRET ratio was measured for at least 10 min using a microplate reader after the addition of furimazine and indicated concentration of compounds.

### Cell permeabilization and calculation of intracellular NMN

HEK 293T cells that stably express NMoRI were subjected to permeabilization in the presence of 15 μM digitonin and varying concentrations of NMN for 15 min at room temperature. Subsequently, the permeabilization solution was removed by aspiration and replaced with 100 μL of HEPES buffer (20 mM HEPES, pH 7.2, and 140 mM NaCl) that include furimazine at 1000-fold dilution, 1 mM ATP, and known NMN concentrations. The pH of the HEPES buffer was adjusted to 8.0 for the estimation of mitochondrial NMN. BRET ratios were recorded using a microplate reader, and intracellular NMN concentrations were determined by referencing standard curves.

### Measurement of cADPR levels in cultured cell

Cell cultures were treated with 500 μM NAM, NMN, NMNH, NR, and NRH for 6h. After incubation, cells were washed with cold PBS and lysed with 0.5 N perchloric acid. The concentration of cADPR was analyzed by cycling assay as described previously.^48^

### NAD^+^ detection by PARP1cd sensor

The construction of PARP1cd NAD^+^ sensor followed established procedures described in previous publication.^30^ To ensure mitochondrial expression of PARP1cd, a COXVIII peptide was incorporated. HEK 293T cells that stably expressed mito-PARP1cd were generated. After Slc25a45 knockdown using shRNAs, the cells were washed with PBS and lysed using RIPA buffer supplemented with protease and phosphate inhibitors, as well as 1 mM 3-aminobenzamide. A total of 20 μg of lysate was separated by electrophoresis on a 4-20% precast gel and subsequently subjected to immunoblotting. The involved the use of anti-PAR (Millipore, MABE1031) or anti-Actin antibodies, followed by incubation with secondary antibodies for detection.

### Quantification and statistical analysis

All experimental results are expressed as Mean ± SD. Data comparisons among the groups were performed by using unpaired Student’s t-test or ANOVA analysis. A P value smaller than 0.05 was considered statistically significant.

**Correspondence and requests for materials** should be address to Q.Y.

## ACKNOWLEDGEMENT

This work is supported by National Key R&D Program of China (2021YFF1200300), National Natural Science Foundation of China (22207118, 22307135), Research Projects on Key Areas of General Colleges and Universities, Department of Education of Guangdong Province (2022ZDZX2071), Shenzhen Science and Technology Program (JCYJ20220818100804009, ZDSYS20210623091810032), Natural Science Foundation of Guangdong Province (2023A1515010715), Shenzhen Institute of Advanced Technology, Chinese Academy of Sciences.

## AUTHOR CONTRIBUTIONS

Q.Y., K.L., and L.C. conceived this study. L.C., P.W., G.H., and W.C. executed the experiments. All authors contributed to data analysis or manuscript writing.

## COMPETING INTERESTS

L.C. and Q.Y. are inventors of two patent applications filed by SIAT. Patent numbers are CN202211334667.6 and PCT/CN2022/138161.

