## Supplementary Material for "Quantitative dynamics of intracellular NMN by genetically encoded biosensor"

**Affiliations:**

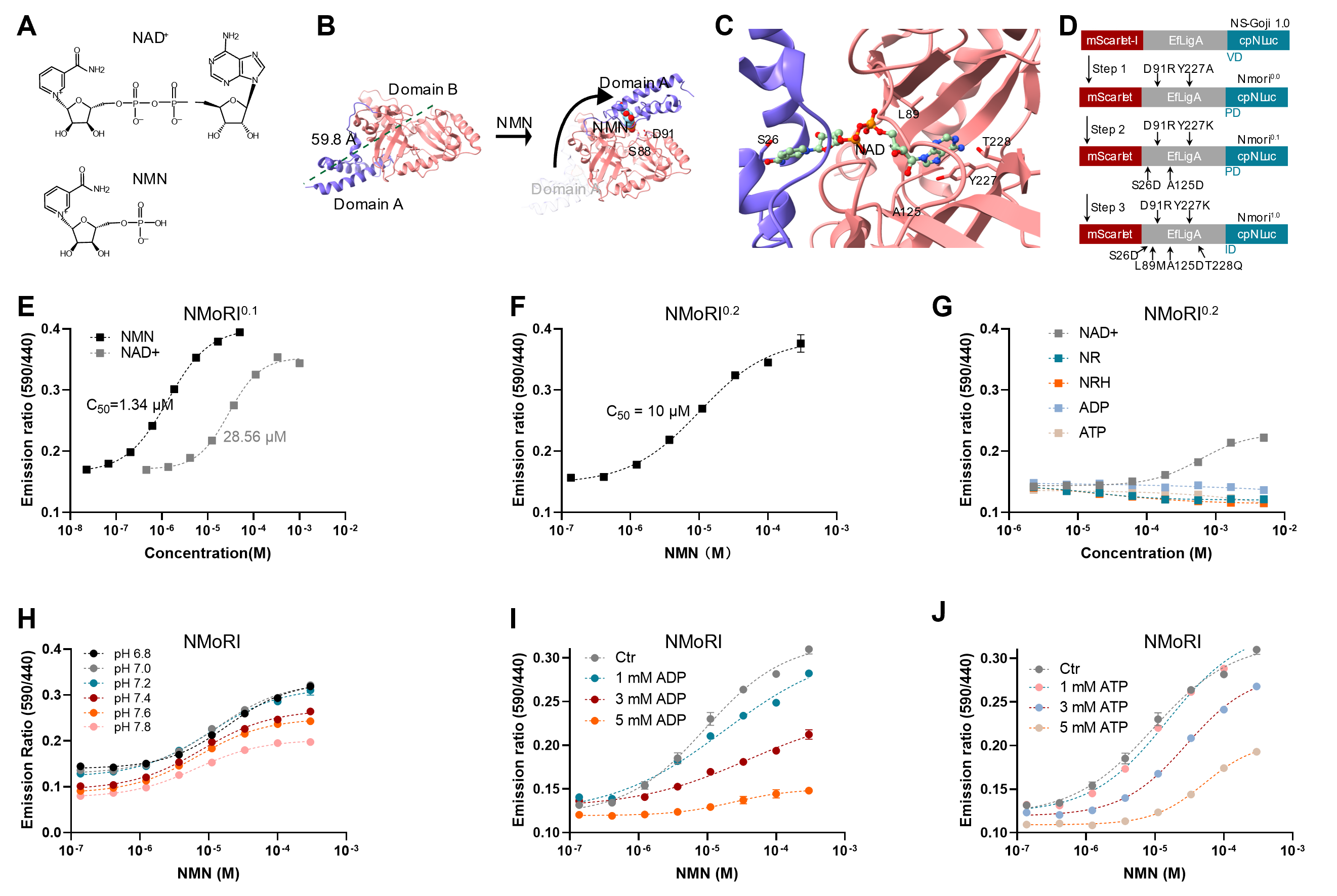

**Figure S1.** **NMoRI sensor development.** (A) Chemical structure of NAD^+^ and NMN. (B) Conceived conformational change of *Ef*LigA variant induced by NMN (PDB code: 1ta8 and 1tae). (C) *Ef*LigA ligand binding pocket (PDB code: 1tae). (D) Schematic showing the workflow of NMN sensor development. (E) Titration curve of NMoRI^0.1^ against NMN and NAD^+^. (F) Titration curve of NMoRI^0.2^ against NMN. (G) Titration curve of NMoRI^0.2^ against NAD^+^, NR, NRH, ADP and ATP. (H) Titration curve of NMoRI at pH 6.8-7.8. (I) Titration curve of NMoRI against NMN with presence of indicated concentrations of ADP. (J) Titration curve of NMoRI against NMN with presence of indicated concentrations of ATP. Data are shown as mean ± SD from at least three independent measurements.

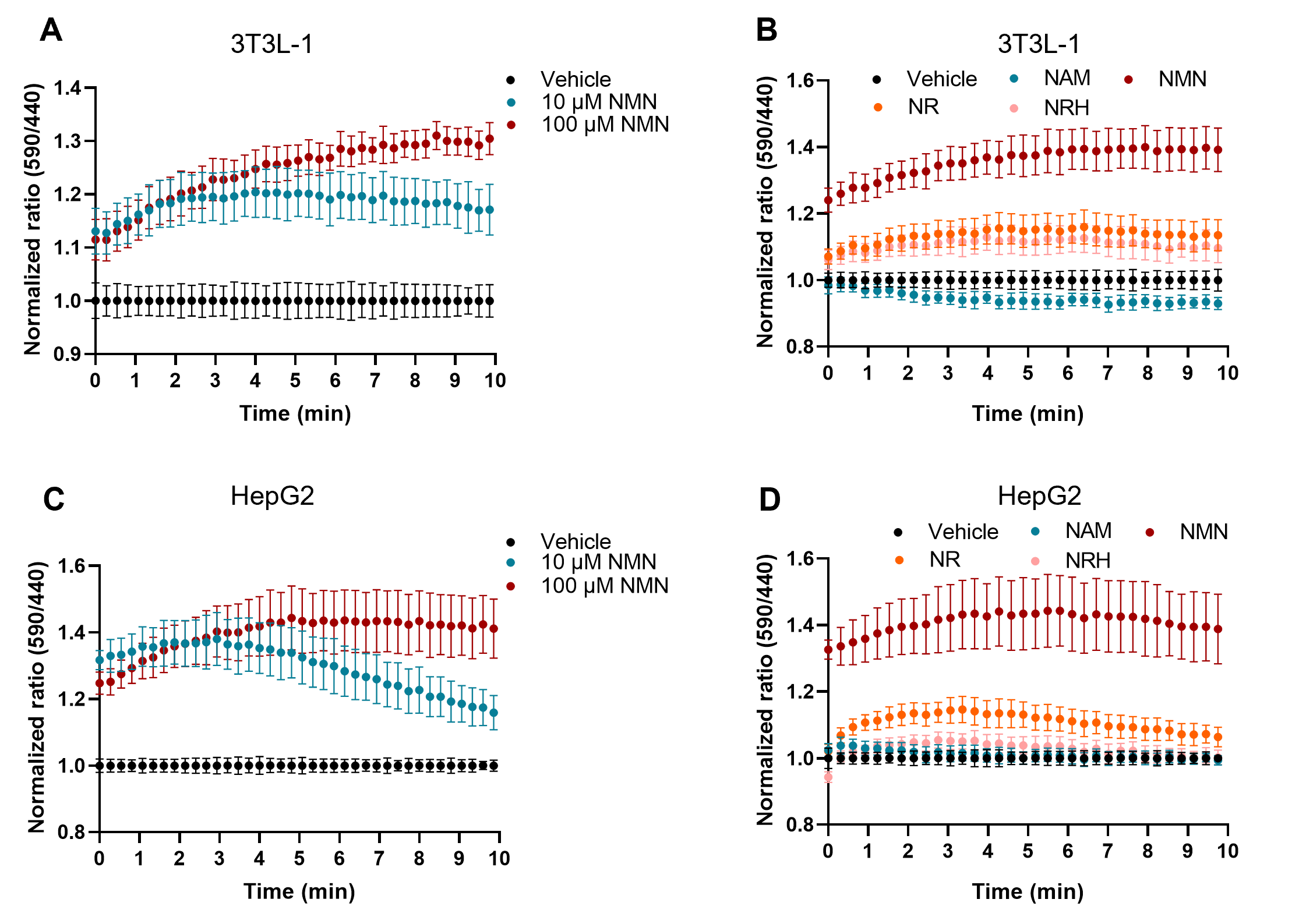
**Figure S2.** **Uptake dynamics of NMN and derivatives in 3T3-L1 and HepG2 cells.** (A) NMoRI BRET ratios of 3T3-L1 cells treated with 0, 10 and 100 μM NMN. (B) NMoRI BRET ratios of 3T3-L1 cells treated with 100 μM NAM, NMN, NR, and NRH. (C) NMoRI BRET ratios of HepG2 cells treated with 0 μM-100 μM NMN. (D) NMoRI BRET ratios of HepG2 cells treated with 100 μM NAM, NMN, NR, and NRH. All values were given as mean ± SD from at least six independent samples.

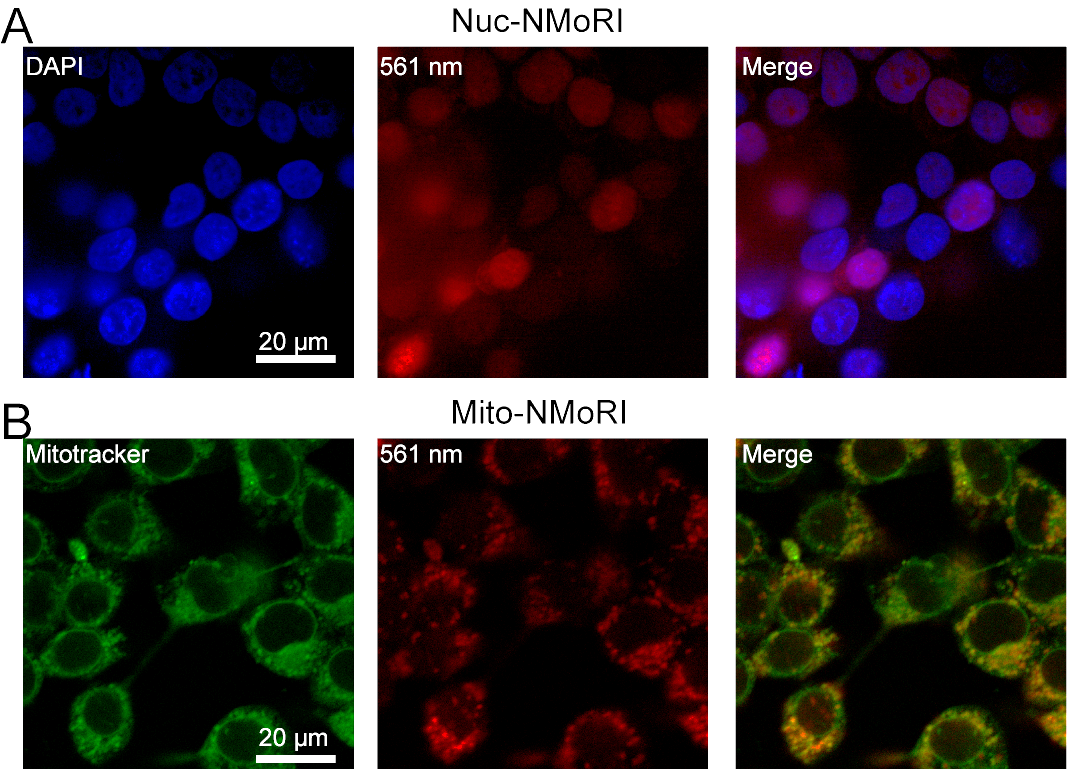
**Figure S3.** **Representative microscopic images of HEK 293T cells expressing NMoRI in nucleus and mitochondria.** (A) NMoRI specifically expressed in nucleus. (B) NMoRI specifically expressed in mitochondria.

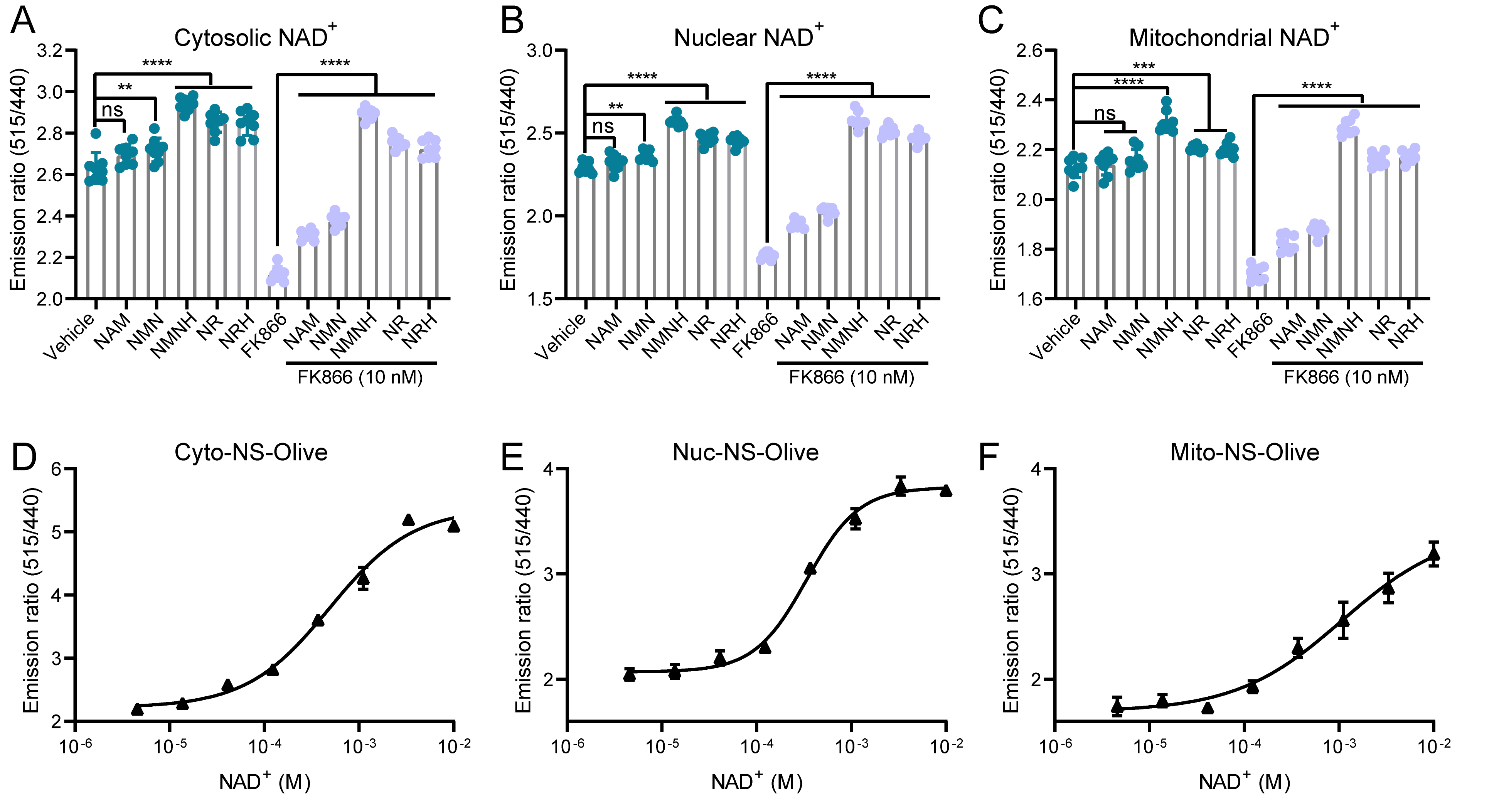
**Figure S4.** **NS-Olive reports compartmentalized NAD^+^ metabolism in HEK 293T cells.** Effects of NAD^+^ precursors (500 μM) on cytosolic (A), nuclear (B), and mitochondrial (C) NAD+ levels with and without FK866 (10 nM), n ≥ 6. Calibration curves for HEK 293T cells expressing NS-Olive in cytosol (D), nucleus (E), and mitochondria (F). Values were given as mean ± SD. Statistical analysis was performed by one-way ANOVA analysis. ns, *P* > 0.05; *, *P* < 0.05; **, *P* < 0.01; ***, *P* < 0.001; ****, *P* < 0.0001.

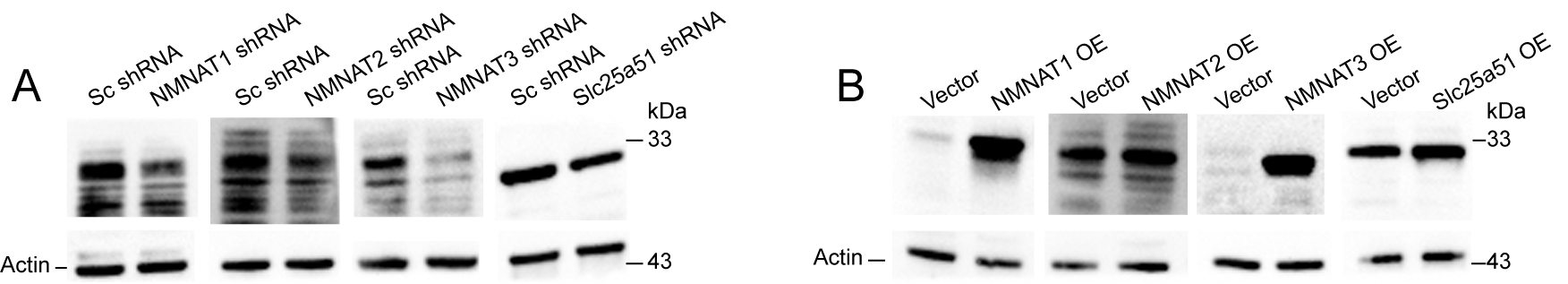

**Figure S5. Western blot analysis of NMoRI expressing HEK 293T cells.** Representative western blot analysis of whole cell lysates from selected genes knockdown (A) or over expression (B) in NMoRI expressing HEK 293T cells.

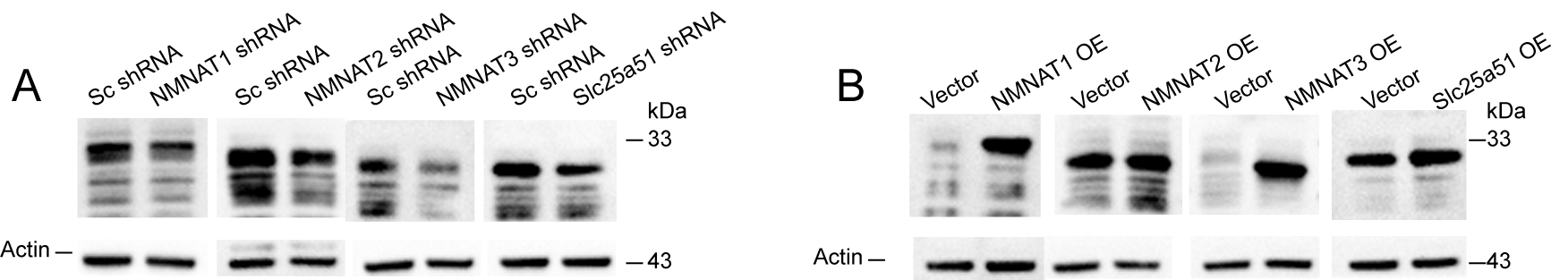

**Figure S6. Western blot analysis of NS-Olive expressing HEK 293T cells.** Representative western blot analysis of whole cell lysates from selected genes knockdown (A) or over expression (B) in NS-Olive expressing HEK 293T cells.

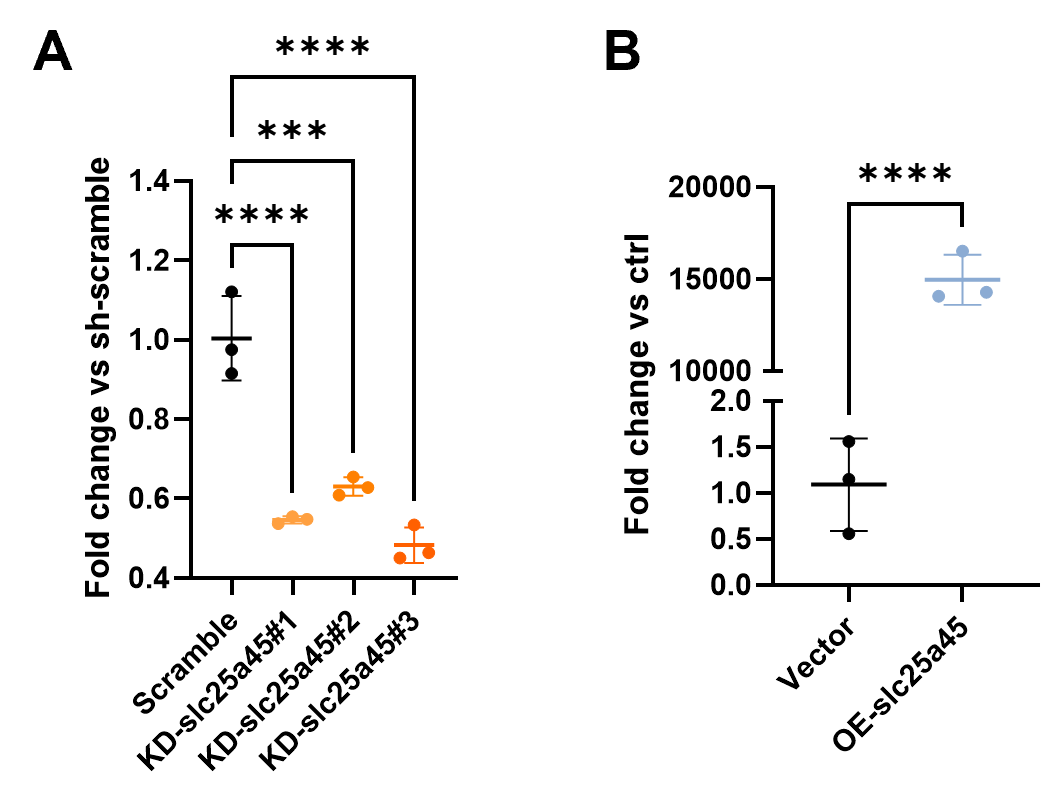
**Figure S7.** **Validation of Slc25a45 knock down and over expression in HEK 293T cells.** Relative gene transcript levels after Slc25a45 knock down (A) and over expression (B). Error bars represent SD. Statistical analysis was performed by one-way ANOVA analysis. ns, *P* > 0.05; *, *P* < 0.05; **, *P* < 0.01; ***, *P* < 0.001; ****, *P* < 0.0001.

**Table S1. Statistics of crystallography data collection and refinement for *EfLigA* variant used in NMoRI.**

| **Data collection** |  |
| --- | --- |
| Space group | P3121 |
| Cell dimensions |  |
| a, b, c (Å) | 92.48, 92.48, 74.29 |
| α, β, γ (°) | 90.00, 90.00, 120.00 |
| Wavelength (Å) | 0.979 |
| Resolution (Å) ^a^ | 30.27-3.00 (3.08-3.00) |
| R_merge_ ^a^ | 0.130 (1.603) |
| I/σ (I) ^a^ | 25.3(3.2) |
| No. of unique observations | 7642(574) |
| No. of total observations | 147067(11561) |
| Completeness (%)^a^ | 99.9 (100.0) |
| CC1/2 | 0.999 (0.927) |
| Redundancy | 19.2 (20.1) |
| **Refinement** |  |
| R_work_/R_free_ | 0.21/0.25 |
| B-factors (Å^2^) |  |
| Protein | 85.85 |
| Ligand | 84.81 |
| Water | 67.51 |
| RMSD |  |
| Bond lengths (Å) | 0.0069 |
| Bond angles (°) | 1.184 |
| Ramachandran statistics (%) |  |
| Favoured | 93.55 |
| Outliers | 0.32 |
| PDB entry | 8JYD |

^a^Data for the highest resolution shell are shown in parentheses.
